# Structural basis for postfusion-specific binding to Respiratory Syncytial Virus F protein by the canonical antigenic site I antibody 131-2a

**DOI:** 10.1101/2025.01.04.631317

**Authors:** Weiwei Peng, Marta Šiborová, Xuesheng Wu, Wenjuan Du, Douwe Schulte, Matti F. Pronker, Cornelis A. M. de Haan, Joost Snijder

## Abstract

The Respiratory Syncytial Virus (RSV) Fusion (F) protein is a major target of antiviral antibodies following natural infection or vaccination and responsible for mediating fusion between the viral envelope and the host membrane. The fusion process is driven by a large-scale conformational change in F, switching irreversibly from the metastable prefusion state to the stable postfusion conformation. Previous research has identified six distinct antigenic sites in RSV-F, termed sites Ø, I, II, III, IV, and V. Of these, only antigenic site I is fully specific to the postfusion conformation of F. A monoclonal antibody 131-2a that specifically targets postfusion F has been widely used as a research tool to probe for postfusion F and to define antigenic site I in serological studies, yet its sequence and precise epitope have remained unknown. Here we use mass spectrometry-based *de novo* sequencing of 131-2a to reverse engineer a recombinant product and study the epitope to define antigenic site I with molecular detail, revealing the structural basis for the antibody’s specificity towards postfusion RSV-F.

## Introduction

Respiratory Syncytial Virus (RSV) is the second major cause for hospital admissions of young infants globally, following malaria. It is estimated that RSV is responsible for approximately 60,000 childhood deaths each year, especially in low-resource settings^1,2^. RSV is a member of the *Pneumoviridae* family. It consists of a negative sense, single stranded RNA genome enveloped in a lipid bilayer containing three virally encoded envelope proteins: the Small Hydrophobic Protein (SHP), Fusion protein (F), and Glycoprotein (G)^3^. Both G and F are involved in host cell attachment and receptor binding and as the name suggests, F also mediates cell entry by fusing the viral envelope with the host membrane, thereby delivering the genetic material of the virus into its host cell^4^. The antibodies that target G and F play a pivotal role in the antiviral immune response and F in particular has become the focus of vaccines and monoclonal antibody therapies that have been recently approved or are currently under clinical development^5–7^.

F is a trimeric class I viral fusion protein that exists in a metastable prefusion conformation. Its conformational change into the stable postfusion conformation drives fusion of the viral envelope with the host membrane. Owing to the drastic conformational changes between the pre- and postfusion states of F, each conformation presents distinct epitopes. Previous studies have described this complex antigenic landscape by defining six antigenic sites: Ø, I, II, III, IV, and V (see Figure 1)^11^. Antigenic sites Ø and V are specific to the prefusion state of F, and while antibodies directed against antigenic site III can bind F in both conformations, they also bind stronger to the prefusion conformation. Antigenic sites II and IV are shared between pre- and postfusion states, but antibodies directed to antigenic site I are fully specific to the postfusion conformation^8^.

**Figure 1.**
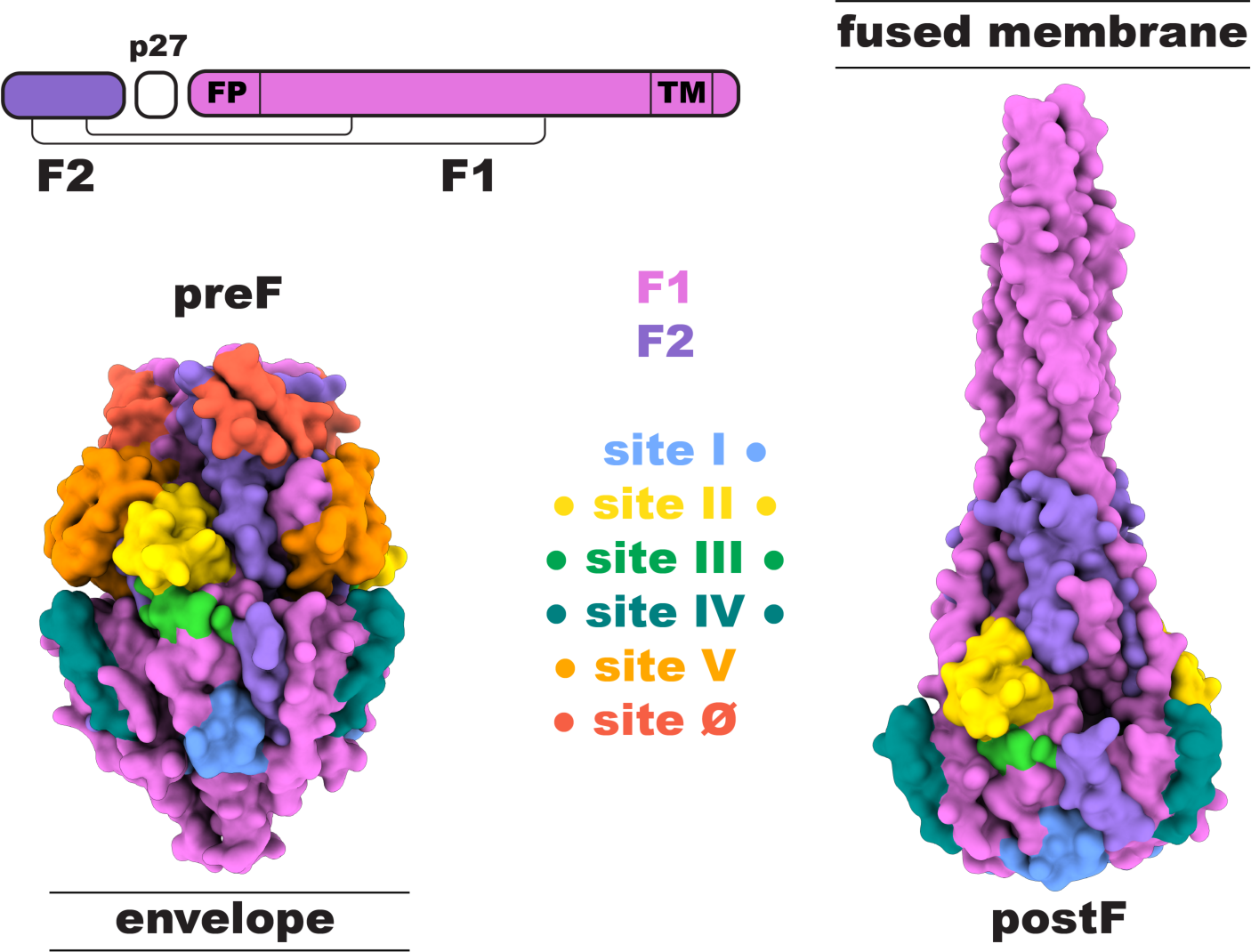
Antigenic sites on the RSV-F protein. The mature RSV-F antigen is proteolytically processed from the F0 precursor into a disulfide-linked F1 and F2 subunit, releasing the p27 peptide. FP: fusion peptide, TM: transmembrane region. (left) Prefusion conformation of RSV-F (PDB ID: 4MMU^8^). (right) Postfusion conformation of RSV-F (PDB ID: 3RRR^9^). Antigenic sites are plotted on the surface with the indicated colors. Specificity of the antigenic sites for pre/postF is indicated with the colored dots. Antigenic site definition follows Hause et al. 2017^10^. The monoclonal antibody 131-2a is widely used to define antigenic site I.

Even in newly produced virions from an infected host cell, F is present on the envelope in a complex mixture of pre- and postfusion states. Meanwhile, neutralizing activity in immune sera can be largely attributed to prefusion F-specific antibodies. The postfusion F on virions has been speculated to act as a decoy to the immune system, directing it towards non-neutralizing epitopes at the expense of effectively neutralizing epitopes present in prefusion F. The antigenic site-I directed antibodies thereby constitute an important, possibly counterproductive component of the antibody response to natural RSV infection or vaccination^12^.

Serological studies have used monoclonal antibody standards to probe the antigenic site-specific antibody response in human subjects. This is typically done in competition binding experiments of the monoclonal antibody standards with polyclonal sera to pre- and postfusion F^13,14^. The mouse monoclonal antibody 131-2a has become the canonical antigenic site I-defining antibody in these studies and is also widely used to specifically detect postfusion F in viral cultures and vaccine preparations by ELISA, Western blot, immunofluorescence microscopy, and cell sorting experiments. While 131-2a was discovered in the early 1980s and has since been widely used in RSV studies, its sequence is not publicly available^15^. Moreover, the epitope of 131-2a on postfusion F has not been studied at a detailed structural level, so the basis of its specificity to postfusion F has not been conclusively established. We have recently developed a mass spectrometry-based workflow to sequence antibodies straight from the purified protein^16–18^, which we apply here to reverse engineer a functional recombinant 131-2a monoclonal antibody. This enabled detailed structural studies of the 131-2a interaction with F by single particle cryogenic electron microscopy (cryoEM), revealing the structural basis for its binding specificity to the postfusion conformation.

## Results

### De novo sequencing 131-2a by mass spectrometry-based bottom-up proteomics

The purified mouse monoclonal antibody 131-2a was sequenced by mass spectrometry, using a bottom-up proteomics approach. The antibody was digested with a panel of 7 proteases in parallel (trypsin, chymotrypsin, α-lytic protease, thermolysin, elastase, lysC and lysN) to generate overlapping peptides for the LC-MS/MS analysis, using an in-solution digestion protocol. Peptides were sequenced from MS/MS spectra, following a hybrid fragmentation scheme with both stepped high-energy collision dissociation (sHCD) and electron-transfer high energy collision dissociation (EThcD) on all peptide precursors. The peptide sequences were predicted from the MS/MS spectra using PEAKS and assembled into the full length heavy and light chain sequences using the in-house developed software Stitch.

This resulted in the identification of a mouse IgG2a antibody with an IGHV1S29 heavy chain, paired with an IGKV3-2 light chain (the full sequence is provided in the Supplementary Information). The depth of coverage for the complementarity determining regions (CDRs) varies from around 10 to 200, indicating a high sequence accuracy (see Supplementary Figure S1). Examples of MS2 spectra supporting the CDRs of both heavy chain and light chain are shown in Figure 2. The heavy and light chains exhibit a typical, moderate degree of somatic hypermutation of an estimated 12% and 5%, respectively. This includes several inferred mutations in the framework regions of the chains.

**Figure 2.**
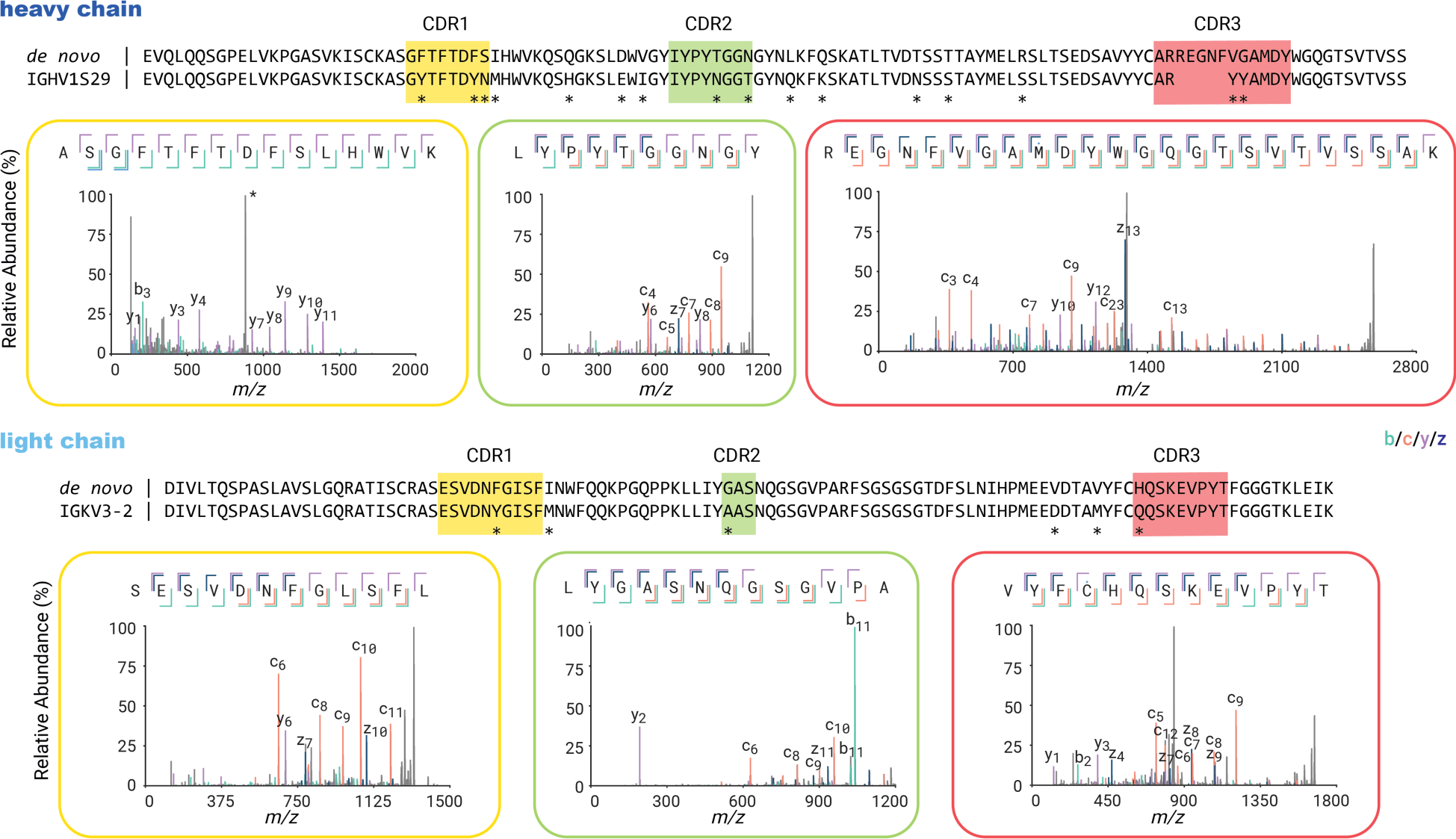
De novo sequencing of 131-2a by mass spectrometry-based bottom-up proteomics. The variable region alignment to the inferred germline sequence is shown for both heavy and light chains. Positions with putative somatic hypermutation are highlighted with asterisks (*). The MS2 spectra supporting the CDR regions are shown beneath the sequence alignment.

### Reverse engineering a functional 131-2a monoclonal antibody

The experimentally determined sequences of 131-2a were reverse translated to DNA with codon optimization for expression in HEK293 cells. The synthetic DNA for the variable domains was inserted into the mouse IgG2a backbone with a C-terminal His_8_-tag on the heavy chain for purification, and the mouse Ig Kappa backbone for the light chain. Plasmids for heavy and light chain were co-transfected in HEK293E cells, with the recombinant 131-2a yielding 95 mg from a 1 L culture after His-tag purification. The reverse engineered 131-2a was then compared with input material for sequencing in Western blot and enzyme-linked immunosorbent assay (ELISA). As shown in Figure 3, the reverse engineered 131-2a binds specifically to postF in a manner that is indistinguishable from the input material. This demonstrates that the mass spectrometry derived sequence yielded a functionally equivalent antibody product.

**Figure 3.**
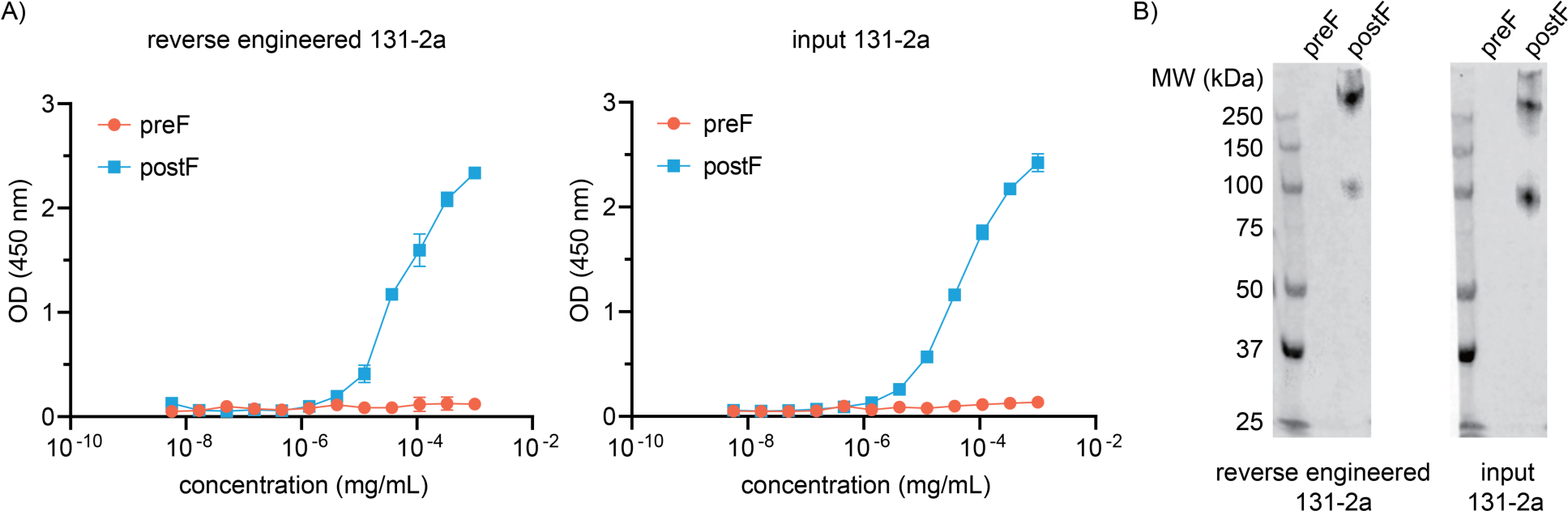
Validation of recombinant 131-2a expressed with mass spectrometry-derived sequence. A) ELISA of pre/postF comparing the sequenced input and reverse engineered 131-2a. B) Western blot analysis following non-denaturing PAGE of pre/postF comparing the sequenced input and reverse engineered 131-2a.

### Epitope mapping of 131-2a

Despite its use as an antibody standard to define antigenic site I and specifically detect the F protein in a postfusion state, the molecular basis of 131-2a’s postfusion specificity is not well understood. Reverse engineering 131-2a enabled us to map the epitope in greater detail using single particle cryoEM (see Supplementary Figure S2). Our reconstructions of the F:131-2a complex recovered a clearly identifiable postF head domain, while the stalk remained largely unresolved. The imaged particles consist of a mixture of three F:131-2a stoichiometries, containing 3:0, 3:1, and 3:2 subunits of each component. The 3:1 complex was most populated in this dataset and refined to a resolution of 3.2 Å. The final map at 3.1 Å resolution reported here is based on C3 reconstructions with symmetry relaxation of the 3:1 and 3:2 complexes, followed by symmetry expansion and combining all poses with a bound Fab into a final local refinement with the constant domains of the Fabs masked out (see Supplementary Table S1).

The epitope of 131-2a spans a single protomer of the postF trimer, burying 1308 Å^2^ of its accessible surface area. Interactions are mediated by all six CDRs of both the heavy and light chain variable domains (see Figure 4). The epitope is a composite of three discontinuous regions from the F1 and F2 subunits, spanning residues 31-42 (F2), 323-332 (F1), and 379-399 (F1). Full lists of buried residues and hydrogen bonds are available in Supplementary Tables S2 and S3.

**Figure 4.**
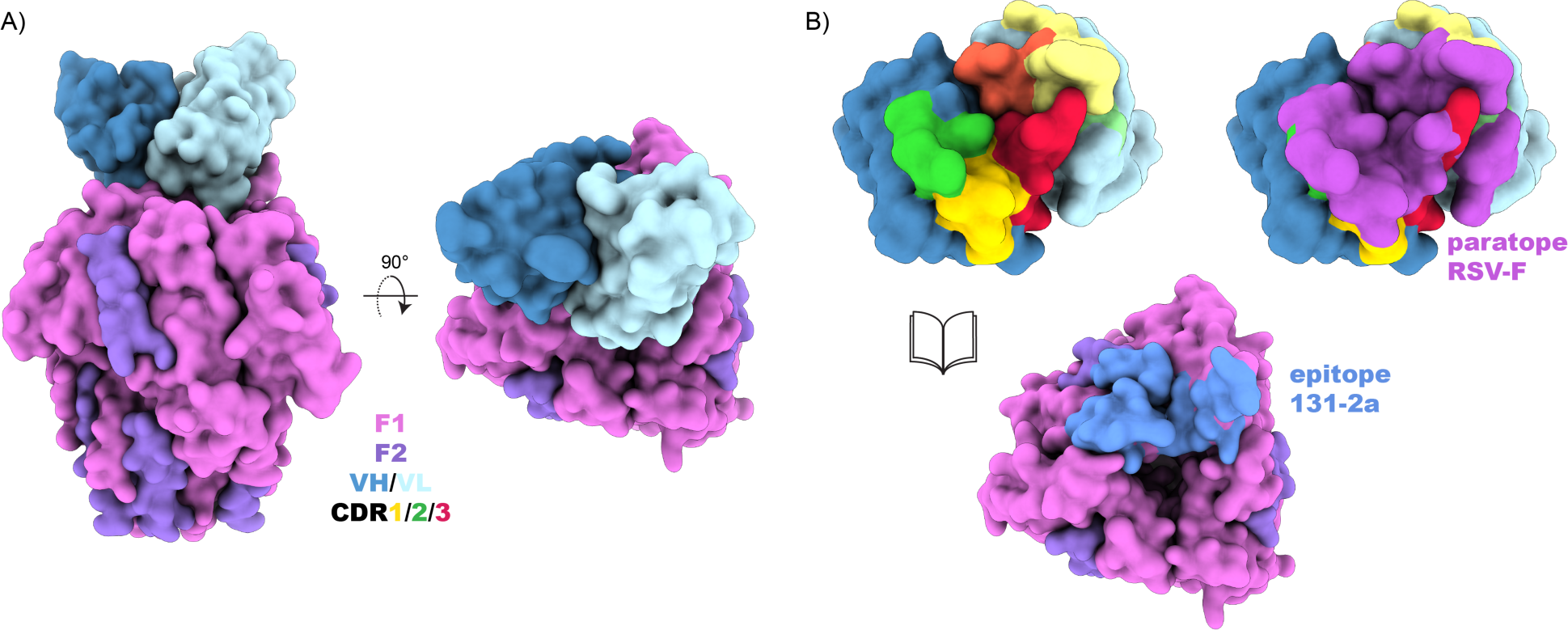
Atomic model of RSV postF bound to the 131-2a Fab. A) Surface representation of postF bound to a single copy of 131-2a’s VH/VL domains. B) Detailed view of the buried surface between 131-2a (paratope) and postF (epitope). CDRs are displayed with the indicated colors using dark/light tones for VH/VL, respectively.

The most extensive contacts are made within residues 379-399 of the F1 subunit (see Figure 5); a looped region exposed on the surface of the postF apex, facing the internal cavity of the trimer’s head domain and stabilized by an internal disulfide bridge (C383-C393). It includes P389, which is a site of known escape mutants to 131-2a and sometimes used to infer binding to antigenic site I by other monoclonal antibodies^9,11^. The peptide bond between P389 and K390 makes a hydrogen bond with the Y55 side chain in the heavy chain framework region 2 (directly flanking CDRH2). The same Y55 residue of the heavy chain is also hydrogen bonded to the side chain of F1-K390, which in turn is also hydrogen bonded to the backbone of N65 in CDRH2. Residue F1- D385 appears to be another critical part of the epitope by hydrogen bonding to both the Y57 side chain of CDRH2 and the backbone of N110 in CDRH3. While the light chain also makes extensive contacts with the 379-399 region, they appear to be exclusively Van der Waals interactions.

**Figure 5.**
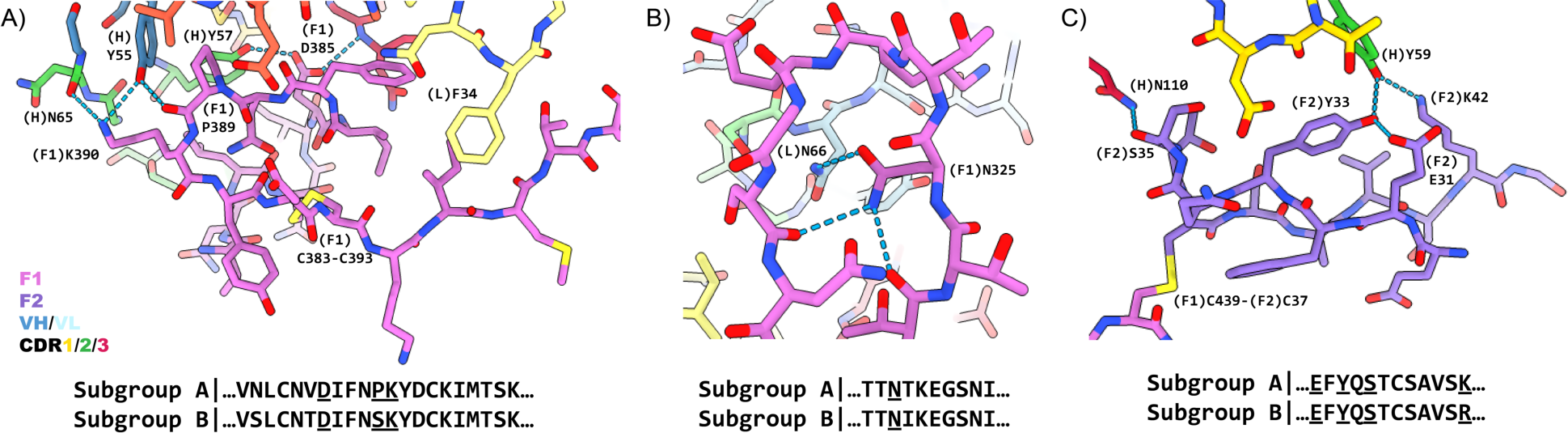
Detailed interactions in the 131-2a epitope on postF. A) Interactions with residues 379-399 of the F1 subunit. B) Interactions with residues 323-332 (C322-C333 loop) in the F1 subunit. C) Interactions with residues 31-42 of the F2 subunit. Key residue numbers with associated chains as indicated, hydrogen bonds are indicated with dashed blue lines. The RSV-F sequence is highly conserved within subgroups A and B, with the corresponding consensus sequences of the epitope displayed. Hydrogen bonded residues to 131-2a are underlined. CDRs are displayed with the indicated colors using dark/light tones for VH/VL, respectively.

Additional contacts are made by the light chain with the loop between the C322-C333 disulfide bridge in the F1 subunits. This loop is disordered in previously published crystal structures of postF^9^, as well as the unbound protomers in the map/model reported here. The loop is stabilized in the interaction with 131-2a by a hydrogen bond between F1-N325 and residue N66 in framework region 3 of the light chain (directly flanking CDRL2). The 131-2a epitope is completed by interactions of the heavy chain with residues 31-42 of the F2 subunit at the apex of the postF head domain. The region spans the disulfide bridge between the F1 and F2 subunits (at F1-C439 and F2-C37). The interaction includes hydrogen bonds between the backbone of F2-S35 with the N110 side chain of CDRH3, as well as F2-Y33 and F2-K42 with Y59 of CDRH2.

The determined model of the 131-2a interaction with postF can be used to rationalize most of the somatic hypermutations observed in the mature VH/VL sequences compared to the inferred germline precursors (see Supplementary Figure S3). The mutated sites within or directly flanking the CDRs of both heavy and light chain are directly involved in the interaction with postF. In addition, we inferred mutations in both the heavy and light chains that are directly situated at the interface of the VH/VL domains, possibly involved in pairing of the chains. Finally, the IGHV1S29 germline of the heavy chain carries a predicted N-linked glycosylation site at N82 of the framework 3 region. A putative glycan at this location is predicted to shield a large portion of the paratope, rationalizing the observed N82T mutation in the mature 131-2a sequence to allow free access to the postF interface in the absence of the N-linked glycan (see Supplementary Figure S3B).

Based on the epitope determined here, the structural basis for 131-2a’s specificity to the postF conformation can be explained (see Figure 6). While the conformation of the epitope itself remains stable between preF and postF (see Supplementary Figure S4), access to the F1-379-399 region is sterically blocked by the C-terminus of the F1 subunit in the preF state. This region contains the α10 helix of the heptad repeat B (HRB) region, which refolds to the helical stalk domain of the postF state. The specificity of 131-2a to postF is therefore negatively determined by steric blocking of the epitope in the preF conformation. This mechanism is broadly similar to the postfusion specificity of another site I directed antibody, ADI-14359^19^, though it differs in some crucial details of how the epitope is blocked in the preF state (see Supplementary Figure S5). While 131-2a binds at the apex of the postF head domain in a rather sharp angle relative to the symmetry axis of the trimer, the angle of approach is more obtuse, and the epitope more peripheral, for ADI-14359. Contacts by ADI-14359 are mediated for the largest part by the heavy chain, but even though the light chain has a minor contribution binding to postF, it has a critical role in mediating the postF specificity. The epitope of ADI-14359 is still fully accessible in the preF conformation and not blocked by the α10 helix of the F1-HRB region. Rather, the extended C-terminus of the F1 subunit, directly preceding the α10 helix, augments the antiparallel beta sheet formed by the looped 31-42 region of F2, which is shared in both 131-2a and ADI-14359 epitopes, and produces a steric clash with the ADI-14359 light chain in the preF conformation.

**Figure 6.**
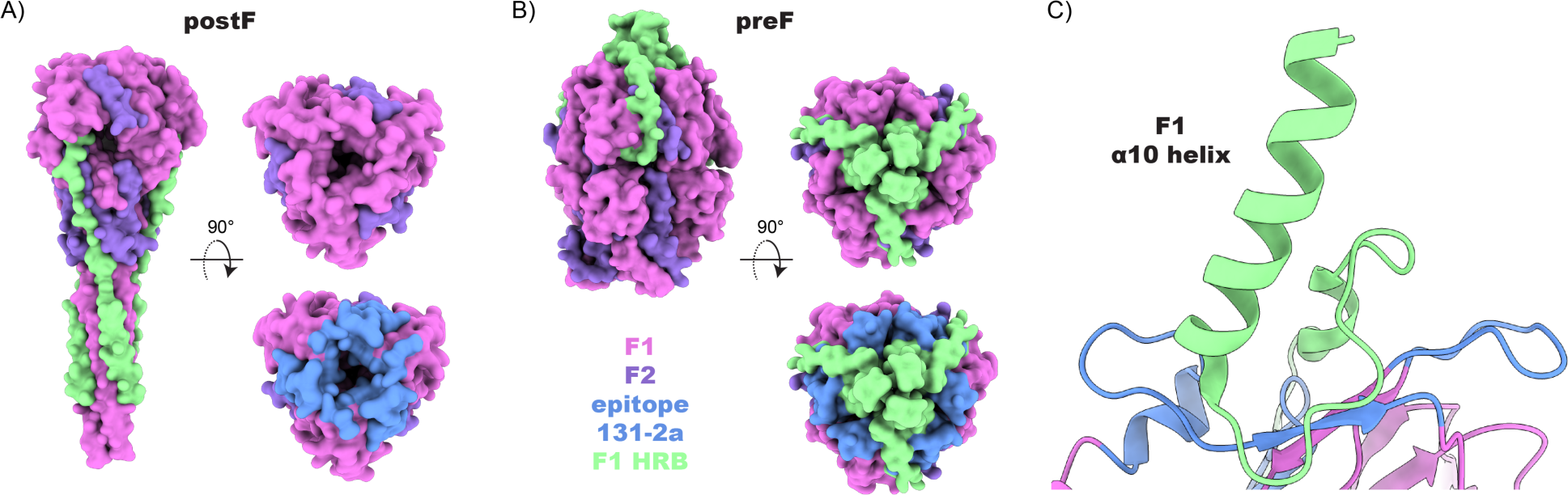
Structural basis for postF specificity of 131-2a. A) Surface representation of postF crystal structure (PDB ID: 3RRR). B) Surface representation of preF crystal structure (PDB ID: 4MMU). C) Cartoon representation of single preF protomer at the 131-2a epitope. While the conformation of the 131-2a epitope remains the same in preF vs postF, access to the epitope is blocked in preF by the α10 helix and heptad repeat B (HRB) of the F1 C-terminus, which refolds into the helical stalk region of postF.

Curiously, the structure of postF bound to ADI-14359, like 131-2a, was also solved in a 3:1 stoichiometry. While we can’t exclude that in both cases the substoichiometric binding is a sample preparation or processing artefact, it is interesting to note that the 131-2a epitope determined here would predict a very close fit for the fully occupied 3:3 complex (see Supplementary Figure S6). The light chains of the three 131-2a variable domains point inwards over the central cavity of the postF trimer and while technically no direct clashes are observed, there is no space for as much as a water molecule between them. This may result in less favorable binding energies for a second and third bound Fab and could go some way to explain the low occupancy observed in our preparations, despite the molar excess of 131-2a Fab added to postF.

## Conclusions

In this work, the application of mass spectrometry-based bottom-up proteomics allowed us to derive the full sequence of the canonical antigenic site-I-defining, anti-RSV-F monoclonal antibody 131-2a. This enabled the reverse engineering of a functional recombinant 131-2a antibody, which was demonstrated to possess equivalent binding specificity to RSV postF when compared to the sequenced input material.

This enabled single particle cryoEM to map the epitope of 131-2a in detail. It revealed a composite epitope encompassing residues 379-399 in the F1 subunit, with additional stabilizing contacts to the loop spanning C322-C333 in F1, and extensive contacts with residues 31-42 in F2. While the conformation of the epitope remains similar in the prefusion conformation of F, access is hindered by the C-terminus of the F1 subunit, which refolds to the stalk region in the postfusion conformation. The postF specificity of 131-2a can thus be explained by negative selection of binding in the prefusion conformation. *De novo* sequencing of 131-2a by mass spectrometry enabled an in-depth molecular characterization of antigenic site I of the RSV-F protein, shedding new light on decades of serological studies characterizing the antibody response to RSV infection and vaccination.

## Methods

### Expression and Purification of RSV-F Proteins

Design, expression and purification of RSV pre-fusion F (DSCav1-T4fd, Genbank JX015498) has been described previously^14,20^. Briefly, cDNA encoding pre-fusion RSV F (DSCav1-T4fd) was cloned into the pCD5 expression vector in frame with the CD5 signal peptide coding sequence, followed by sequences encoding a C-terminal T4 fibritin trimerization motif, thrombin site, and StrepII-tag (IBA, Germany). Pre-fusion F was expressed transiently in HEK-293T cells [ATCC, CRL-11268] and secreted protein was purified from culture supernatants using Strep-tactin Sepharose beads (IBA) following the manufacturer’s protocol and as described previously^14,21^. RSV post-fusion F corresponds to Flys-GCN described previously except that the StrepII-tag was not present in the current protein and that it was expressed in CHO cells^21^. This construct was generated by cloning cDNAs encoding the RSV F ectodomain (amino acids 26 to 515, Genbank: JX015498.1) in frame with a CD5 signal peptide-encoding sequence and followed by sequences coding for the GCN4 isoleucine zipper trimerization motif and a a LysM peptidoglycan binding domain^22,23^. In addition, arginines in the two furin-cleavage sites were substituted by lysines. This protein was previously used in a clinical trial^24^ and did not display D25 reactivity after its prolonged storage at -80°C.

### MS-based sequencing of 131-2a

*Sample preparation* – 24 μg of 131-2a (MAB8599P-K, Sigma) was denatured and reduced in 2% sodium deoxycholate (SDC), 200 mM Tris-HCl, 10 mM tris(2-carboxyethyl)phosphine (TCEP), 40 mM iodoacetic acid, pH 8.5 at 95 °C for 10 min, followed with 20 min incubation at room temperature in the dark for alkylation. 3 μg sample was then digested by one of the following proteases trypsin, chymotrypsin, α-lytic protease, thermolysin, elastase, lysC and lysN in a 1:50 ratio (w:w) in a total volume of 100 uL of 50 mM ammonium bicarbonate at 37 °C overnight. After digestion, SDC was removed by adding 2 µL formic acid (FA) and centrifugation at 14000 ×g for 20 min. Following centrifugation, the supernatant containing the peptides was collected for desalting on a 30 µm Oasis HLB 96-well plate (Waters). The Oasis HLB sorbent was activated with 100% acetonitrile and subsequently equilibrated with 10% formic acid in water. Next, peptides were bound to the sorbent, washed twice with 10% formic acid in water and eluted with 100 µL of 50% acetonitrile/5% formic acid in water (v/v). The eluted peptides were vacuum-dried and reconstituted in 100 µL 2% FA.

*Mass Spectrometry* –The digested peptides (single injection of 0.2 ug) were separated by online reversed phase chromatography on an Agilent 1290 UHPLC (column packed with Poroshell 120 EC C18; dimensions 50 cm x 75 µm, 2.7 µm, Agilent Technologies) coupled to a Thermo Scientific Orbitrap Fusion mass spectrometer. Samples were eluted over a 90 min gradient from 0% to 35% acetonitrile at a flow rate of 0.3 μL/min. Peptides were analyzed with a resolution setting of 60000 in MS1. MS1 scans were obtained with standard AGC target, maximum injection time of 50 ms, and scan range 350-2000. The precursors were selected with a 3 m/z window and fragmented by stepped HCD as well as EThcD. The stepped HCD fragmentation included steps of 25%, 35% and 50% NCE. EThcD fragmentation was performed with calibrated charge-dependent ETD parameters and 27% NCE supplemental activation. For both fragmentation types, MS2 scan were acquired at 30000 resolution, 800% Normalized AGC target, 250 ms maximum injection time, scan range 120-3500.

*Data analysis* – MS/MS spectra were used to determine *de novo* peptide sequences using PEAKS Studio (version 11.5)^25^. We used a tolerance of 20 ppm and 0.02 Da for MS1 and MS2, respectively. Carboxymethylation was set as fixed modification of cysteine. Oxidation of methionine, tryptophan, and histidine, and pyroglutamic acid modification of N-terminal glutamic acid and glutamine were set as additional variable modifications. The CSV file containing all the *de novo* sequenced peptides was exported for further analysis. Stitch (version 1.5) was used for the template-based assembly^26,27^. The mouse antibody database from IMGT was used as templates^28,29^. The cutoff score for the *de novo* sequenced peptide was set as 85 and the cutoff score for the template matching was set as 16. A first iteration was performed with automatic V-J segment joining, which failed for the heavy chain due to minimal overlap between the overhanging reads. The CDRH3 region was constructed manually from the overhanging peptides assembled at the ends of the V and J segments, followed by a second iteration of Stitch, assembling the input reads to the first iteration draft sequences of the full heavy and light chains. The resulting consensus sequence was manually corrected to include the conserved cysteine residues at the start of CDRH1 and CDRH3. The determined variable domain sequences were plugged into the identified constant domain sequences from Uniprot to arrive at the final sequences reported here. V-genes and CDR boundaries were assigned with the IMGT DomainGapAlign webserver (https://www.imgt.org/3Dstructure-DB/cgi/DomainGapAlign.cgi).

### Cloning and Expression of recombinant 131-2a

To recombinantly express full-length 131-2a, the proteomic sequences of both the light and heavy chains were reverse-translated and codon-optimized for expression in human cells using the Thermo Fisher webtool (https://www.thermofisher.com/order/gene-design/index.html). For the linker and Fc region of the heavy chain, the standard mouse IgG2A amino acid sequence (IMGT database) was used. An N-terminal secretion signal peptide derived from human IgG light chain (MEAPAQLLFLLLLWLPDTTG) was added to the N-termini of both heavy and light chains. BamHI and NotI restriction sites were added to the 5′ and 3′ ends of the coding regions, respectively. Only for the light chain, a double stop codon was introduced at the 3′ site before the NotI restriction site. The coding regions were subcloned using BamHI and NotI restriction-ligation into a pRK5 expression vector with a C-terminal octahistidine tag between the NotI site and a double stop codon 3′ of the insert, so that only the heavy chain has a C-terminal AAAHHHHHHHH sequence for nickel-affinity purification (the triple alanine resulting from the NotI site). After the sequence was validated by Sanger Sequencing, the HC/LC were mixed in a 1:1 DNA ratio and expressed in HEK293 cells by the ImmunoPrecise Antibodies (Europe) B.V company. After expression the culture supernatant of the cells was harvested and purified using a HisPur Ni-NTA Resin (Thermo Fisher Scientific). After purification, 131-2a was buffer exchanged and concentrated into phosphate-buffered saline (PBS) using a centrifugal filter (Amicon). The expression plasmids described above are available through Addgene (https://www.addgene.org/Joost_Snijder/).

### 131-2a Fab generation

The full 131-2a IgG was digested by immobilized papain (ThermoFisher) in digestion buffer (0.22 mM Cysteine HCl, 20 mM phosphate buffer, pH 7) at 37 °C, 1000 rpm shaking for 5 hours. After digestion, the Fc segment was removed by incubation with protein A agarose Resin (Thermo Fisher Scientific) at room temperature for 15 minutes. The 131-2a Fab was further purified by size-exclusion chromatography using a Superdex 200 Increase 10/300 GL column (Cytiva) equilibrated in PBS buffer.

### Western blot

Binding of 131-2a was analyzed by western blot assay utilizing both prefusion and postfusion proteins mentioned above^14,20,21^. Briefly, 0.25 μg of either post- or pre-fusion RSV F proteins were mixed with native protein buffer (Bio-Rad, 1610738) and then loaded onto a 7% polyacrylamide gel devoid of SDS, accompanied by protein standards (Bio-Rad, 1610375). The proteins were transferred to a cellulose nitrate membrane using a Trans-Blot Turbo Transfer System (Bio-Rad). Following this, the membrane was subjected to blocking utilizing 3% BSA alongside 0.1% Tween 20. Horseradish peroxidase (HRP)-conjugated rabbit anti-mouse IgG (Dako, P0260) was used at a dilution of 1:5,000 for detection of the 131-2a antibody. Visualization was performed using BeyoECL Moon (Beyotime). The Western blots were scanned using an imaging system (Odyssey).

### ELISA

Nunc MaxiSorp ELISA plates (Thermo Fisher Scientific) were coated with 50 ng of RSV F and incubated overnight at 4 °C, followed by three washing steps with PBS containing 0.05% Tween 20. Plates were blocked with 2% bovine serum albumin (BSA; Fitzgerald) in PBS with 0.1% Tween 20 at 4 °C overnight. Subsequently, 131-2a (home-made or commercial (MAB8599P-K, Sigma)) antibodies were allowed to bind the plates at 3-fold serial dilutions, starting at 1 μg/ml diluted in PBS containing 2% BSA and 0.1% Tween 20, at RT for 1 hour. After washing, plates were incubated with 1:1000 diluted HRP–conjugated rabbit anti-mouse IgG (Dako, P0260) for 1 hour at RT. HRP reactivity with tetramethylbenzidine substrate (BioFX) was determined by measuring absorbtion at 450 nm using an ELISA plate reader (EL-808, BioTek).

### CryoEM sample preparation

RSV postF and 131-2a Fabs were mixed in 3:4 molar ratio and incubated 30 minutes on ice. Sample was diluted to 0.2 mg/ml in PBS and was pipetted onto a holey carbon-coated copper grid (R1.2/1.3, mesh 200; Quantifoil), blotted and vitrified by plunging into liquid ethane using an FEI Vitrobot Mark IV (Thermo Fisher Scientific).

### CryoEM data collection

The vitrified sample was transferred to a Titan Krios electron microscope (Thermo Fisher Scientific) operated under cryogenic conditions and at an acceleration voltage of 300 kV. A total of 4330 micrographs were collected at a magnification of 105,000× on a K3 direct electron detector in counted super-resolution mode, resulting in a calibrated pixel size of 0.418 Å/pix. Imaging was done under low-dose conditions (total dose 50 e^−^/Å^2^) and defocus values ranging from −0.8 to −2.0 μm. The 2.52 second exposure was fractionated into 50 frames and saved in tiff format. Automated data acquisitions were performed using the software EPU with AFIS (Thermo Fisher Scientific).

### Cryo-EM data processing

Data was processed in Cryosparc in successive versions from v4.2.1 to v4.5.3^30^. The movies were motion-corrected and 2x binned using PatchMotionCorrection. CTF parameters were estimated with the PatchCTF function using default parameters. Micrographs were filtered to an estimated max. resolution from PatchCTF of <5 Å, selecting 3802 for the final processed dataset.

An initial set of 2.1M particles was picked with the Blobpicker function, extracted using an original boxsize of 360px and binned 4x for further processing. Following two successive rounds of 2D classification, 563k particles were selected for Ab Initio reconstruction into 8 classes. The resulting volumes could be grouped as RSV-F trimers with 0, 1 or 2 copies of the bound Fab, along with a smaller unidentified class, somewhat resembling a free Fab. Particles corresponding to postF trimers with either 0, 1, or 2 Fabs were re-extracted with a 360px box size binned 1.5x, resulting in maps of 3.4 Å, 3.3 Å and 3.4 Å resolution, respectively, following non-uniform refinement.

The postF +1Fab map was used to generate templates for the template picker function, resulting in 2.4M new picks. Using the previously reconstructed maps of postF +0, +1 or +2 Fabs as starting volumes, along with the ab initio class of the smaller Fab-like class, the full stack of 2.4M particles (original boxsize 360px binned 4x) was used as input for an initial round of heterorefinement. The particles for each assigned volume were then subjected to two additional successive rounds of heterorefinement with the same 4 starting volumes selecting only the particles for the target class for each new round. This resulted in 186k particles classified as postF +0 Fab, 200k particles +1 Fab, and 97k particles +2 Fabs.

After extraction of the classified particles with original boxsize 360px binned 1.5x, the particles were reconstructed to 3.1, 3.2 and 3.3 Å resolution for +0, +1 +2 Fabs, respectively, following successive non-uniform and local refinements (C1 symmetry). The maps for postF with +1 and +2 Fabs were then reconstructed with C3 symmetry using symmetry relaxation (marginalization), the maps aligned, symmetry expanded, and the poses with Fabs bound combined into the final reconstruction at 3.1 Å resolution following local refinement with the constant domains of the Fab masked out.

### Cryo-EM model building and refinement

Software used in this project was curated by SBGrid^31^. As a starting point for the postF portion of the model, a homology model of our RSV-F construct based on the published postF crystal structure (PDB ID: 3RRR^9^) prepared using the SWISS-MODEL webserver, was rigid body fitted using the ‘fit in map’ function of ChimeraX (v1.8)^32,33^. The VH/VL structure of 131-2a was predicted with the SAbPred ABodyBuilder2 webserver, adopting IMGT numberings for the chains^34^. The predicted VH/VL structure was rigid body fitted in ChimeraX and combined into the same model with the postF trimer. Density for the loop spanning the C322-C333 disulfide bridge of the F1 subunit was only visible in the 131-2a bound protomer of the postF trimer, which was manually fitted using Coot (WinCoot v0.9.8.1)^35^. The resulting model was used as input for flexible fitting using the Namdinator webserver with default parameters^36^. Regions outside the map (termini, C322-C333 loops of the two unbound postF protomers, and the HRA/HRB helical stalk region) were pruned from the model, followed by a final round of flexible fitting in Namdinator with default settings. One final round of refinement in Phenix (v1.21.2.5419) using non-bonded weight of 500 resulted in the final model reported here^37^. Interfaces were analyzed using the PISA webserver^38^. All figures were prepared using ChimeraX (v1.8).

The model of glycosylated 131-2a T82N was prepared by structure prediction with the SAbPred ABodyBuilder2 webserver, followed by grafting of an A1F glycan with the GLYCOSHIELD webserver^39^.

## Data Availability

The raw LC-MS/MS files and analyses have been deposited to the ProteomeXchange Consortium via the PRIDE partner repository with the dataset identifier PXD059427. The final reported map has been deposited in EMDB under ID 52444. The additional maps of RSV-F with 0, 1, and 2 Fabs bound are deposited in EMDB under IDs 52446, 52447, and 52448, respectively. The atomic model of postF with bound VH/VL of 131-2a has been deposited to the Protein Data Bank with the accession code 9HVW. The plasmids for 131-2a expression in mammalian cells are made available through Addgene, under plasmid ID 214965 and 215331 for the heavy and light chains, respectively.

## Acknowledgements

We would like to thank Daniel Hurdiss (Utrecht University) and his team for fruitful discussions regarding the EM data processing and model building and Bert Jan Haijema (3D-PharmXchange) for providing details on the RSV F protein used in this study. This research was funded by the Dutch Research Council NWO Gravitation 2013 BOO, Institute for Chemical Immunology (ICI; 024.002.009) and the European Research Council Executive Agency HORIZON ERC-2022-STG (FLAVIR; 101077640) to J.S. We additionally acknowledge support from NWO, funding the MS and EM facilities through NEMI (184.034.014) and the X-omics (184.034.019) Road Map programs.

## References

(1) Cohen, C.; Zar, H. J. Deaths from RSV in Young Infants—the Hidden Community Burden. The Lancet Global Health 2022, 10 (2), e169–e170. 10.1016/S2214-109X(21)00558-1.

(2) Reichert, H.; Suh, M.; Jiang, X.; Movva, N.; Bylsma, L. C.; Fryzek, J. P.; Nelson, C. B. Mortality Associated With Respiratory Syncytial Virus, Bronchiolitis, and Influenza Among Infants in the United States: A Birth Cohort Study From 1999 to 2018. The Journal of Infectious Diseases 2022, 226 (Supplement_2), S246–S254. 10.1093/infdis/jiac127.

(3) Bergeron, H. C.; Tripp, R. A. Immunopathology of RSV: An Updated Review. Viruses 2021, 13 (12), 2478. 10.3390/v13122478.

(4) Battles, M. B.; McLellan, J. S. Respiratory Syncytial Virus Entry and How to Block It. Nat Rev Microbiol 2019, 17 (4), 233–245. 10.1038/s41579-019-0149-x.

(5) Higgins, D.; Trujillo, C.; Keech, C. Advances in RSV Vaccine Research and Development – A Global Agenda. Vaccine 2016, 34 (26), 2870–2875. 10.1016/j.vaccine.2016.03.109.

(6) Haynes, L. M. Progress and Challenges in RSV Prophylaxis and Vaccine Development. The Journal of Infectious Diseases 2013, 208 (suppl_3), S177–S183. 10.1093/infdis/jit512.

(7) Shaw, C. A.; Ciarlet, M.; Cooper, B. W.; Dionigi, L.; Keith, P.; O’Brien, K. B.; Rafie-Kolpin, M.; Dormitzer, P. R. The Path to an RSV Vaccine. Current Opinion in Virology 2013, 3 (3), 332–342. 10.1016/j.coviro.2013.05.003.

(8) Mousa, J. J.; Sauer, M. F.; Sevy, A. M.; Finn, J. A.; Bates, J. T.; Alvarado, G.; King, H. G.; Loerinc, L. B.; Fong, R. H.; Doranz, B. J.; Correia, B. E.; Kalyuzhniy, O.; Wen, X.; Jardetzky, T. S.; Schief, W. R.; Ohi, M. D.; Meiler, J.; Crowe, J. E. Structural Basis for Nonneutralizing Antibody Competition at Antigenic Site II of the Respiratory Syncytial Virus Fusion Protein. Proceedings of the National Academy of Sciences 2016, 113 (44), E6849–E6858. 10.1073/pnas.1609449113.

(9) McLellan, J. S.; Yang, Y.; Graham, B. S.; Kwong, P. D. Structure of Respiratory Syncytial Virus Fusion Glycoprotein in the Postfusion Conformation Reveals Preservation of Neutralizing Epitopes. Journal of Virology 2011, 85 (15), 7788–7796. 10.1128/jvi.00555-11.

(10) Hause, A. M.; Henke, D. M.; Avadhanula, V.; Shaw, C. A.; Tapia, L. I.; Piedra, P. A. Sequence Variability of the Respiratory Syncytial Virus (RSV) Fusion Gene among Contemporary and Historical Genotypes of RSV/A and RSV/B. PLOS ONE 2017, 12 (4), e0175792. 10.1371/journal.pone.0175792.

(11) López, J. A.; Bustos, R.; Örvell, C.; Berois, M.; Arbiza, J.; García-Barreno, B.; Melero, J. A. Antigenic Structure of Human Respiratory Syncytial Virus Fusion Glycoprotein. Journal of Virology 1998, 72 (8), 6922–6928. 10.1128/jvi.72.8.6922-6928.1998.

(12) Huang, J.; Diaz, D.; Mousa, J. J. Antibody Epitopes of Pneumovirus Fusion Proteins. Front. Immunol. 2019, 10. 10.3389/fimmu.2019.02778.

(13) Anderson, L. J.; Hierholzer, J. C.; Tsou, C.; Hendry, R. M.; Fernie, B. F.; Stone, Y.; McIntosh, K. Antigenic Characterization of Respiratory Syncytial Virus Strains with Monoclonal Antibodies. The Journal of Infectious Diseases 1985, 151 (4), 626–633. 10.1093/infdis/151.4.626.

(14) Widjaja, I.; Wicht, O.; Luytjes, W.; Leenhouts, K.; Rottier, P. J. M.; van Kuppeveld, F. J. M.; Haijema, B. J.; de Haan, C. A. M. Characterization of Epitope-Specific Anti-Respiratory Syncytial Virus (Anti-RSV) Antibody Responses after Natural Infection and after Vaccination with Formalin-Inactivated RSV. Journal of Virology 2016, 90 (13), 5965–5977. 10.1128/jvi.00235-16.

(15) Fernie, B. F.; Cote, P. J.; Gerin, J. L. Classification of Hybridomas to Respiratory Syncytial Virus Glycoproteins. Proceedings of the Society for Experimental Biology and Medicine 1982, 171 (3), 266–271. 10.3181/00379727-171-41509.

(16) Peng, W.; Pronker, M. F.; Snijder, J. Mass Spectrometry-Based De Novo Sequencing of Monoclonal Antibodies Using Multiple Proteases and a Dual Fragmentation Scheme. J. Proteome Res. 2021, 20 (7), 3559–3566. 10.1021/acs.jproteome.1c00169.

(17) Peng, W.; den Boer, M. A.; Tamara, S.; Mokiem, N. J.; van der Lans, S. P. A.; Bondt, A.; Schulte, D.; Haas, P.-J.; Minnema, M. C.; Rooijakkers, S. H. M.; van Zuilen, A. D.; Heck, A. J. R.; Snijder, J. Direct Mass Spectrometry-Based Detection and Antibody Sequencing of Monoclonal Gammopathy of Undetermined Significance from Patient Serum: A Case Study. J. Proteome Res. 2023, 22 (9), 3022–3028. 10.1021/acs.jproteome.3c00330.

(18) Peng, W.; Giesbers, K. C.; Šiborová, M.; Beugelink, J. W.; Pronker, M. F.; Schulte, D.; Hilkens, J.; Janssen, B. J.; Strijbis, K.; Snijder, J. Reverse-Engineering the Anti-MUC1 Antibody 139H2 by Mass Spectrometry–Based de Novo Sequencing. Life Science Alliance 2024, 7 (6). 10.26508/lsa.202302366.

(19) Goodwin, E.; Gilman, M. S. A.; Wrapp, D.; Chen, M.; Ngwuta, J. O.; Moin, S. M.; Bai, P.; Sivasubramanian, A.; Connor, R. I.; Wright, P. F.; Graham, B. S.; McLellan, J. S.; Walker, L. M. Infants Infected with Respiratory Syncytial Virus Generate Potent Neutralizing Antibodies That Lack Somatic Hypermutation. Immunity 2018, 48 (2), 339–349.e5. 10.1016/j.immuni.2018.01.005.

(20) McLellan, J. S.; Chen, M.; Joyce, M. G.; Sastry, M.; Stewart-Jones, G. B. E.; Yang, Y.; Zhang, B.; Chen, L.; Srivatsan, S.; Zheng, A.; Zhou, T.; Graepel, K. W.; Kumar, A.; Moin, S.; Boyington, J. C.; Chuang, G.-Y.; Soto, C.; Baxa, U.; Bakker, A. Q.; Spits, H.; Beaumont, T.; Zheng, Z.; Xia, N.; Ko, S.-Y.; Todd, J.-P.; Rao, S.; Graham, B. S.; Kwong, P. D. Structure-Based Design of a Fusion Glycoprotein Vaccine for Respiratory Syncytial Virus. Science 2013, 342 (6158), 592–598. 10.1126/science.1243283.

(21) Rigter, A.; Widjaja, I.; Versantvoort, H.; Coenjaerts, F. E. J.; Roosmalen, M. van; Leenhouts, K.; Rottier, P. J. M.; Haijema, B. J.; Haan, C. A. M. de. A Protective and Safe Intranasal RSV Vaccine Based on a Recombinant Prefusion-Like Form of the F Protein Bound to Bacterium-Like Particles. PLOS ONE 2013, 8 (8), e71072. 10.1371/journal.pone.0071072.

(22) Harbury, P. B.; Zhang, T.; Kim, P. S.; Alber, T. A Switch Between Two-, Three-, and Four-Stranded Coiled Coils in GCN4 Leucine Zipper Mutants. Science 1993, 262 (5138), 1401–1407. 10.1126/science.8248779.

(23) van Roosmalen, M. L.; Kanninga, R.; El Khattabi, M.; Neef, J.; Audouy, S.; Bosma, T.; Kuipers, A.; Post, E.; Steen, A.; Kok, J.; Buist, G.; Kuipers, O. P.; Robillard, G.; Leenhouts, K. Mucosal Vaccine Delivery of Antigens Tightly Bound to an Adjuvant Particle Made from Food-Grade Bacteria. Methods 2006, 38 (2), 144–149. 10.1016/j.ymeth.2005.09.015.

(24) Ascough, S.; Vlachantoni, I.; Kalyan, M.; Haijema, B.-J.; Wallin-Weber, S.; Dijkstra-Tiekstra, M.; Ahmed, M. S.; van Roosmalen, M.; Grimaldi, R.; Zhang, Q.; Leenhouts, K.; Openshaw, P. J.; Chiu, C. Local and Systemic Immunity against Respiratory Syncytial Virus Induced by a Novel Intranasal Vaccine. A Randomized, Double-Blind, Placebo-Controlled Clinical Trial. Am J Respir Crit Care Med 2019, 200 (4), 481–492. 10.1164/rccm.201810-1921OC.

(25) PEAKS: powerful software for peptide de novo sequencing by tandem mass spectrometry - Ma - 2003 - Rapid Communications in Mass Spectrometry - Wiley Online Library. https://analyticalsciencejournals.onlinelibrary.wiley.com/doi/10.1002/rcm.1196 (accessed 2025-01-02).

(26) Schulte, D.; Peng, W.; Snijder, J. Template-Based Assembly of Proteomic Short Reads For *De Novo* Antibody Sequencing and Repertoire Profiling. Anal. Chem. 2022, 94 (29), 10391–10399. 10.1021/acs.analchem.2c01300.

(27) Schulte, D.; Snijder, J. A Handle on Mass Coincidence Errors in De Novo Sequencing of Antibodies by Bottom-up Proteomics. J. Proteome Res. 2024, 23 (8), 3552–3559. 10.1021/acs.jproteome.4c00188.

(28) Ehrenmann, F.; Kaas, Q.; Lefranc, M.-P. IMGT/3Dstructure-DB and IMGT/DomainGapAlign: A Database and a Tool for Immunoglobulins or Antibodies, T Cell Receptors, MHC, IgSF and MhcSF. Nucleic Acids Research 2010, 38 (suppl_1), D301– D307. 10.1093/nar/gkp946.

(29) Lefranc, M.-P. IMGT® Databases, Web Resources and Tools for Immunoglobulin and T Cell Receptor Sequence Analysis, Http://Imgt.Cines.Fr. Leukemia 2003, 17 (1), 260–266. 10.1038/sj.leu.2402637.

(30) Punjani, A.; Rubinstein, J. L.; Fleet, D. J.; Brubaker, M. A. cryoSPARC: Algorithms for Rapid Unsupervised Cryo-EM Structure Determination. Nat Methods 2017, 14 (3), 290–296. 10.1038/nmeth.4169.

(31) Morin, A.; Eisenbraun, B.; Key, J.; Sanschagrin, P. C.; Timony, M. A.; Ottaviano, M.; Sliz, P. Collaboration Gets the Most out of Software. eLife 2013, 2, e01456. 10.7554/eLife.01456.

(32) Waterhouse, A.; Bertoni, M.; Bienert, S.; Studer, G.; Tauriello, G.; Gumienny, R.; Heer, F. T.; de Beer, T. A. P.; Rempfer, C.; Bordoli, L.; Lepore, R.; Schwede, T. SWISS-MODEL: Homology Modelling of Protein Structures and Complexes. Nucleic Acids Research 2018, 46 (W1), W296–W303. 10.1093/nar/gky427.

(33) UCSF ChimeraX: Tools for structure building and analysis - Meng - 2023 - Protein Science - Wiley Online Library. https://onlinelibrary.wiley.com/doi/10.1002/pro.4792 (accessed 2024-12-18).

(34) Abanades, B.; Wong, W. K.; Boyles, F.; Georges, G.; Bujotzek, A.; Deane, C. M. ImmuneBuilder: Deep-Learning Models for Predicting the Structures of Immune Proteins. Commun Biol 2023, 6 (1), 1–8. 10.1038/s42003-023-04927-7.

(35) Emsley, P.; Lohkamp, B.; Scott, W. G.; Cowtan, K. Features and Development of Coot. Acta Cryst D 2010, 66 (4), 486–501. 10.1107/S0907444910007493.

(36) Kidmose, R. T.; Juhl, J.; Nissen, P.; Boesen, T.; Karlsen, J. L.; Pedersen, B. P. Namdinator – Automatic Molecular Dynamics Flexible Fitting of Structural Models into Cryo-EM and Crystallography Experimental Maps. IUCrJ 2019, 6 (4), 526–531. 10.1107/S2052252519007619.

(37) Liebschner, D.; Afonine, P. V.; Baker, M. L.; Bunkóczi, G.; Chen, V. B.; Croll, T. I.; Hintze, B.; Hung, L.-W.; Jain, S.; McCoy, A. J.; Moriarty, N. W.; Oeffner, R. D.; Poon, B. K.; Prisant, M. G.; Read, R. J.; Richardson, J. S.; Richardson, D. C.; Sammito, M. D.; Sobolev, O. V.; Stockwell, D. H.; Terwilliger, T. C.; Urzhumtsev, A. G.; Videau, L. L.; Williams, C. J.; Adams, P. D. Macromolecular Structure Determination Using X-Rays, Neutrons and Electrons: Recent Developments in Phenix. Acta Cryst D 2019, 75 (10), 861–877. 10.1107/S2059798319011471.

(38) Krissinel, E.; Henrick, K. Inference of Macromolecular Assemblies from Crystalline State. Journal of Molecular Biology 2007, 372 (3), 774–797. 10.1016/j.jmb.2007.05.022.

(39) Tsai, Y.-X.; Chang, N.-E.; Reuter, K.; Chang, H.-T.; Yang, T.-J.; von Bülow, S.; Sehrawat, V.; Zerrouki, N.; Tuffery, M.; Gecht, M.; Grothaus, I. L.; Colombi Ciacchi, L.; Wang, Y.-S.; Hsu, M.-F.; Khoo, K.-H.; Hummer, G.; Hsu, S.-T. D.; Hanus, C.; Sikora, M. Rapid Simulation of Glycoprotein Structures by Grafting and Steric Exclusion of Glycan Conformer Libraries. Cell 2024, 187 (5), 1296–1311.e26. 10.1016/j.cell.2024.01.034.

