## Supplementary Information for "Structural basis for postfusion-specific binding to Respiratory Syncytial Virus F protein by the canonical antigenic site I antibody 131-2a"

>131-2a_heavychain
EVQLQQSGPELVKPGASVKISCKASGFTFTDFSIHWVKQSQGKSLDWVGYIYPYTGGNGYNLK
FQSKATLTVDTSSTTAYMELRSLTSEDSAVYYCARREGNFVGAMDYWGQGTSVTVSSAKTTA
PSVYPLAPVCGDTTGSSVTLGCLVKGYFPEPVTLTWNSGSLSSGVHTFPAVLQSDLYTLSSSV
TVTSSTWPSQSITCNVAHPASSTKVDKKIEPRGPTIKPCPPCKCPAPNLLGGPSVFIFPPKIKDV
LMISLSPIVTCVVVDVSEDDPDVQISWFVNNVEVHTAQTQTHREDYNSTLRVVSALPIQHQDW
MSGKEFKCKVNNKDLPAPIERTISKPKGSVRAPQVYVLPPPEEEMTKKQVTLTCMVTDFMPEDI
YVEWTNNGKTELNYKNTEPVLDSDGSYFMYSKLRVEKKNWVERNSYSCSVVHEGLHNHHTTK
SFSRTPGK

>131-2a_lightchain
DIVLTQSPASLAVSLGQRATISCRASESVDNFGISFINWFQQKPGQPPKLLIYGASNQGSGVPA
RFSGSGSGTDFSLNIHPMEEVDTAVYFCHQSKEVPYTFGGGTKLEIKRADAAPTVSIFPPSSEQ
LTSGGASVVCFLNNFYPKDINVKWKIDGSERQNGVLNSWTDQDSKDSTYSMSSTLTLTKDEYE
RHNSYTCEATHKTSTSPIVKSFNRNCCSNTTGSKT

Supplementary Table S1. CryoEM data collection and refinement statistics.

| **Reconstruction** | **RSV-F postfusion trimer with one 131-2a Fab** |
|---|---|
| **Data collection and processing** | **EMDB-52444/52446/52447/52448** |
| Magnification | 105000 |
| Voltage (kV) | 300 |
| Electron exposure (e-/Å$^2$) | 50 |
| Defocus range (μm) | -0.8 to -2.0 |
| Pixel size (Å) | 0.83 |
| Symmetry imposed | C1/C3/C1/C1 |
| Map resolution (Å) | 3.1/3.1/3.2/3.3 |
| FSC threshold | 0.143 |
| **Refinement** | **PDB ID: 9HVW** |
| Initial model used (PDB ID) | 3RRR |
| Map sharpening *B* factor (Å$^2$) | -105 |
| # non-hydrogen atoms | 8961 |
| # residues | 1149 |
| *B* factor (Å$^2$) | -84 |
| R.m.s. deviations | |
| Bond lengths (Å) | 0.004 |
| Bond angles (°) | 0.817 |
| Validation | |
| MolProbity score | 1.8 |
| Clashscore | 6.17 |
| Poor rotamers (%) | 1.53 |
| Ramachandran plot | |
| Favored (%) | 95.37 |
| Allowed (%) | 4.63 |
| Disallowed (%) | 0 |

Supplementary Table S2. Buried residues in the 131-2a interaction with postF (from PISA).

| RSV-F_F1 (chain E) | RSV-F_F2 (chain B) | 131-2a HC (chain H) | 131-2a LC (chain L) |
|---|---|---|---|
| 321L | 31E | 29T | 31N |
| 323T | 32F | 35T | 34F |
| 325N | 33Y | 36D | 35G |
| 327K | 34Q | 37F | 36I |
| 328E | 35S | 38S | 38F |
| 330S | 36T | 40H | 55Y |
| 332I | 40V | 52W | 65S |
| 334L | 42K | 55Y | 66N |
| 336R | | 57Y | 67Q |
| 377S | | 59Y | 68G |
| 378E | | 62T | 69S |
| 380N | | 64G | 70G |
| 381L | | 65N | 107S |
| 383N | | 66G | 108K |
| 384I | | 107R | 109E |
| 385D | | 108E | 114V |
| 386I | | 109G | 116Y |
| 387F | | 110N | |
| 388N | | 112V | |
| 389P | | | |
| 390K | | | |
| 392D | | | |
| 395I | | | |
| 396M | | | |
| 397T | | | |
| 399K | | | |

| |
|---|
| contact |
| H-bonded residue |

Supplementary Table S3. List of hydrogen bond interaction between 131-2a and postF (from PISA).

| from residue | distance (Å) | to residue |
|---|---|---|
| | **F1-to-HC** | |
| E:LYS 390[ NZ ] | 3.26 | H:TYR 55[ OH ] |
| E:ASN 383[ ND2] | 3.76 | H:TYR 59[ OH ] |
| E:LYS 390[ NZ ] | 3.07 | H:ASN 65[ O ] |
| E:ASP 385[ OD1] | 3.30 | H:ASN 110[ N ] |
| E:ASP 385[ OD2] | 2.97 | H:TYR 57[ OH ] |
| E:PRO 389[ O ] | 2.86 | H:TYR 55[ OH ] |
| | **F2-to-HC** | |
| B:TYR 33[ OH ] | 2.87 | H:TYR 59[ OH ] |
| B:SER 35[ O ] | 2.98 | H:ASN 110[ ND2] |
| B:LYS 42 [ NZ ] | 3.05 | H:TYR 59[ OH ] |
| | **F1-to-LC** | |
| E:ASN 325[ OD1] | 2.99 | L:ASN 66[ ND2] |
| E:PHE 387[ O ] | 3.73 | L:TYR 116[ OH ] |

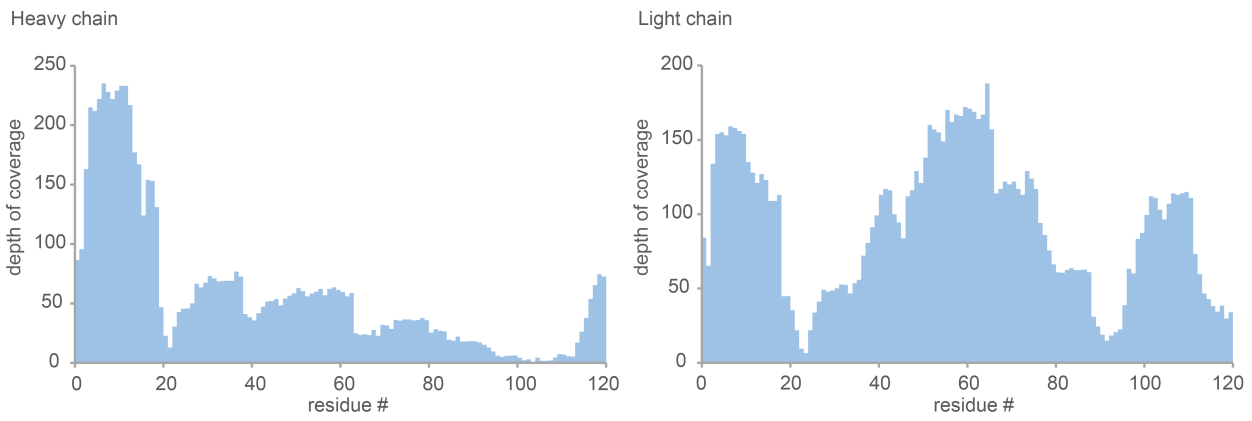

Supplementary Figure S1. Depth of coverage (total number of overlapping peptides mapped per position) for the variable domains of the 131-2a heavy and light chains.

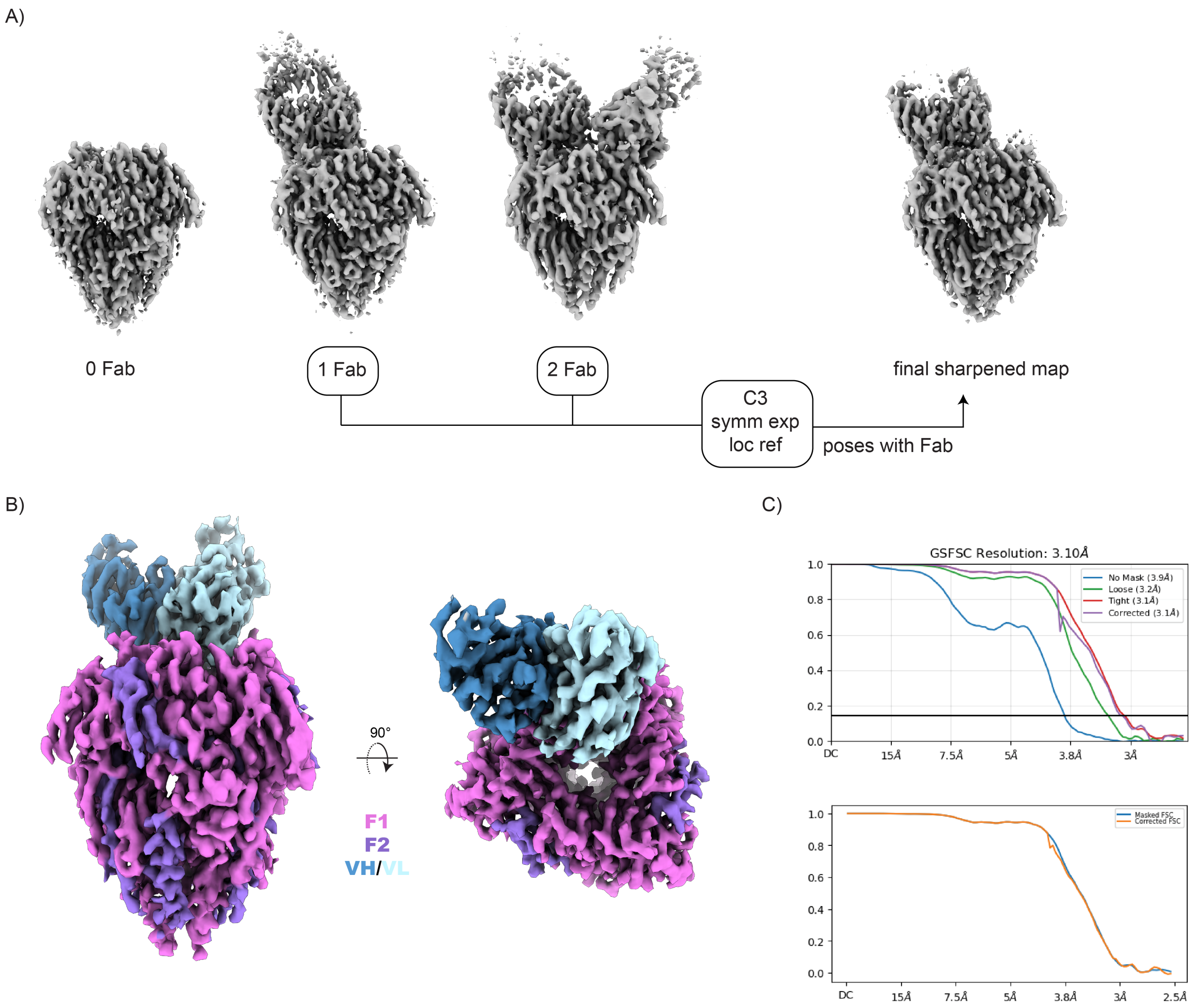

Figure S2. CryoEM single particle reconstruction of postF in complex with 131-2a Fab. A) Final local refinements of 3:0, 3:1 and 3:2 stoichiometries. Following C3 reconstruction with symmetry relaxation (maximization), the 3:1 and 3:2 maps were aligned, symmetry expanded, and the posed with bound Fabs combined in one local refinement to obtain the final map. B) Final sharpened map used for model building. C) FSC plot for the final local refinement (top) and sharpened map (bottom) used for model building.

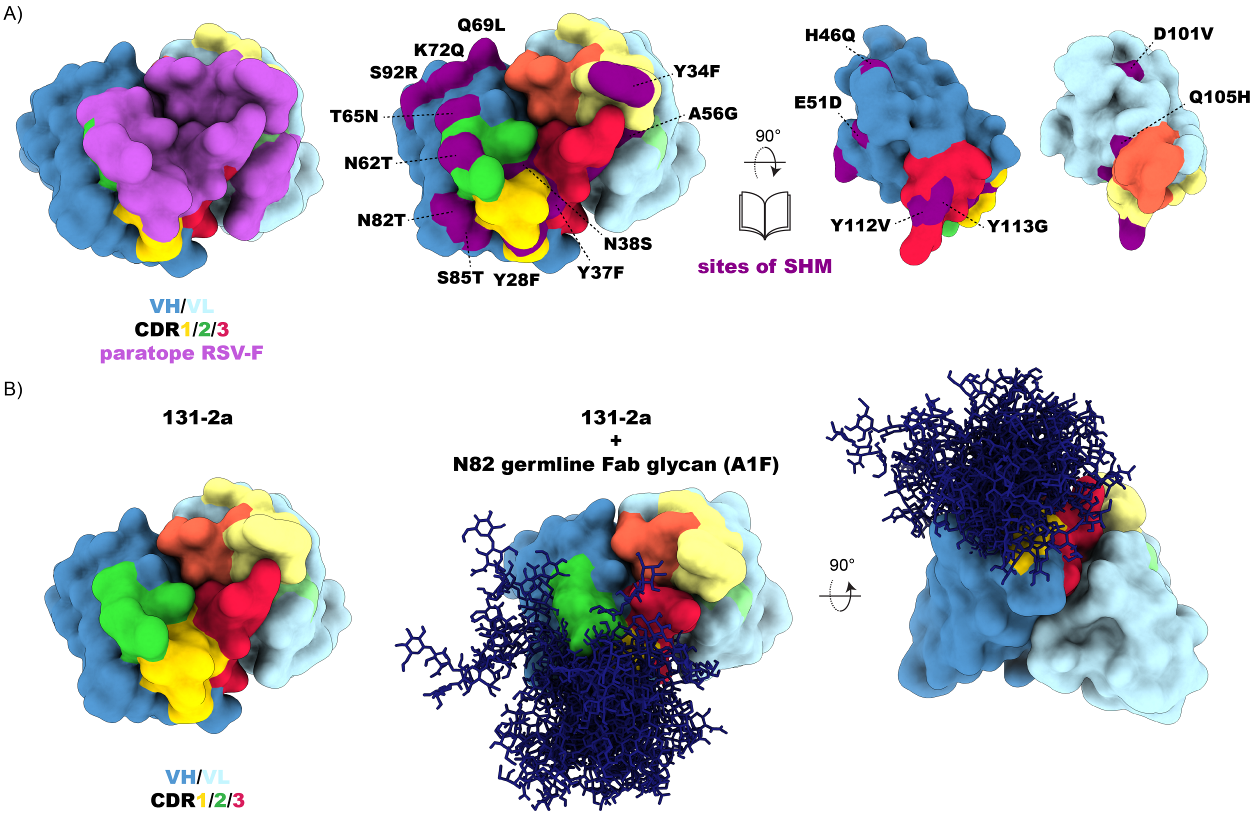

Figure S3. Analysis of inferred somatic hypermutation (SHM) in mature 131-2a sequence. A) Detailed view of paratope (left) compared to sites of SHM (middle), and VH-VL interface (right). B) Model of glycosylated 131-2a without the N82T mutation. An ensemble of 30 glycan conformation was grafted using GLYCOSHIELD, shown as dark blue sticks.

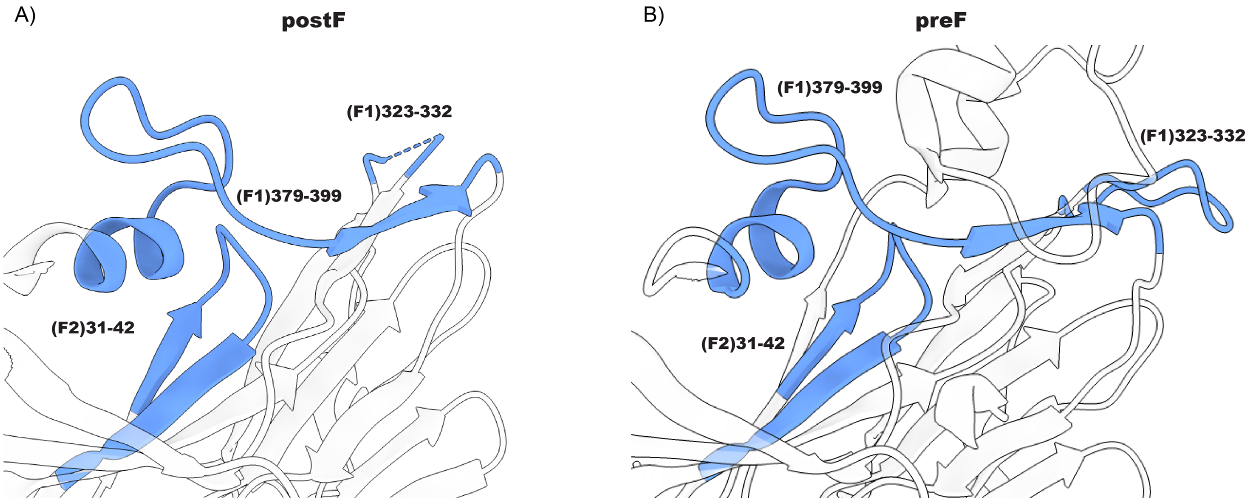

Supplementary Figure S4. Conformation of the 131-2a epitope in postF (A) and preF (B). The epitope is highlighted in blue color. Models used are PDB ID 3RRR and 4MMU for postF and preF, respectively.

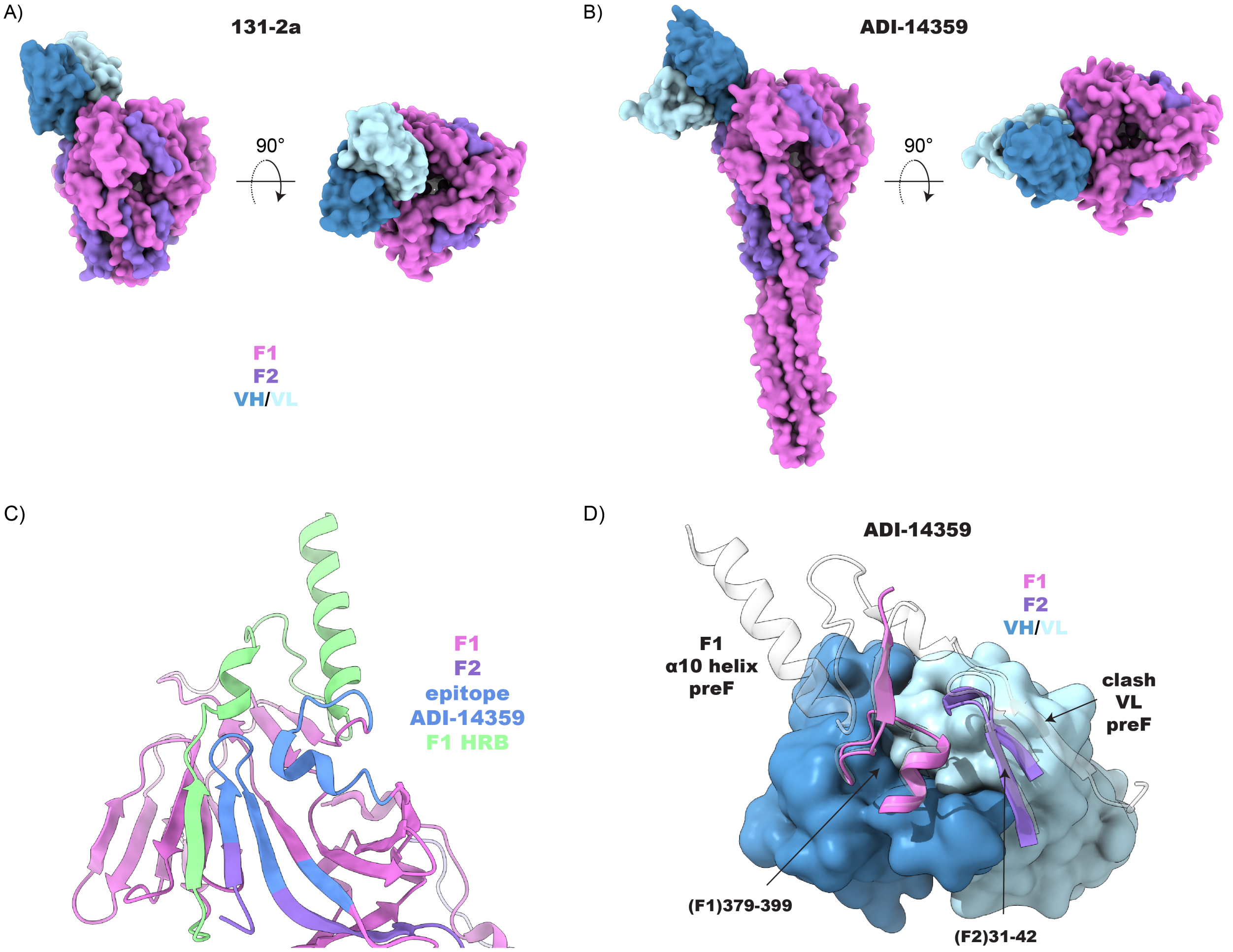

Supplementary Figure S5. Comparison of the 131-2a epitope (A) with the antigenic site I antibody ADI-14359 (PDB ID: 6APB) (B). The cartoon in (C) shows the ADI-14359 epitope mapped onto a single protomer in preF (PDB ID: 4MMU). While the ADI-14359 epitope remains exposed, the extended C-terminus of F1 augments the beta sheet of the F2 region, sterically clashing with the light chain of ADI-14359 (D).

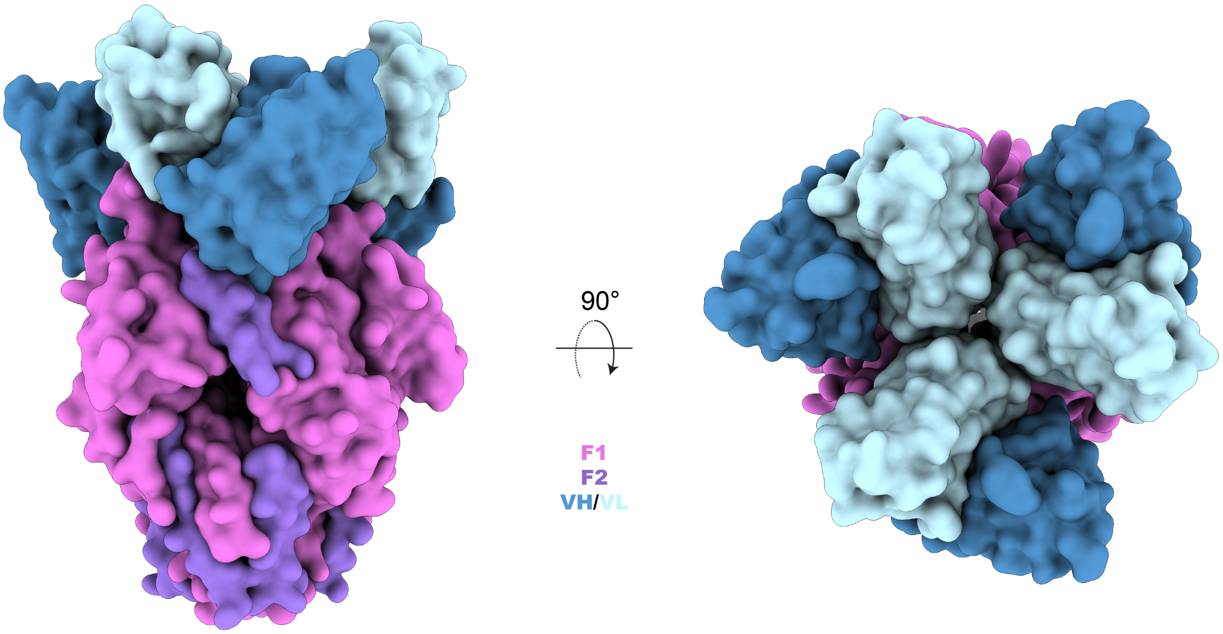

Supplementary Figure S6. Model of postF with 3 copies of 131-2a VH/VL bound. Model was generated by copying the resolved VH/VL domains over to the other protomers in the experimentally determined F:131-2a model. Note the lack of space between the light chains at the central cavity of the postF trimer.
